# Glioblastoma myeloid-derived suppressor cell subsets express differential macrophage migration inhibitory factor receptor profiles that can be targeted to reduce immune suppression

**DOI:** 10.1101/2019.12.19.882555

**Authors:** Tyler J. Alban, Defne Bayik, Balint Otvos, Anja Rabljenovic, Lin Leng, Leu Jia-shiun, Gustavo Roversi, Adam Lauko, Arbaz Momin, Alireza M. Mohammadi, David M. Peereboom, Manmeet S. Ahluwalia, Kazuko Matsuda, Kyuson Yun, Richard Bucala, Michael A. Vogelbaum, Justin D. Lathia

## Abstract

The application of tumor immunotherapy to glioblastoma (GBM) is limited by an unprecedented degree of immune suppression due to factors that include high numbers of immune suppressive myeloid cells, the blood brain barrier, and T cell sequestration to the bone marrow. We previously identified an increase in immune suppressive myeloid-derived suppressor cells (MDSCs) in GBM patients, which correlated with poor prognosis and was dependent on macrophage migration inhibitory factor (MIF). Here we examine the MIF signaling axis in detail in murine MDSC models, GBM-educated MDSCs and human GBM. We found that the monocytic subset of MDSCs (M-MDSCs), expressed high levels of the MIF cognate receptor CD74 and was localized in the tumor microenvironment. In contrast, granulocytic MDSCs (G-MDSCs) expressed high levels of the MIF non-cognate receptor CXCR2 and showed minimal accumulation in the tumor microenvironment. Furthermore, targeting M-MDSCs with ibudilast, a brain penetrant MIF-CD74 interaction inhibitor, reduced MDSC function and enhanced CD8 T cell activity in the tumor microenvironment. These findings demonstrate the MDSC subsets differentially express MIF receptors and may be leveraged for specific MDSC targeting.

## Introduction

Glioblastoma (GBM) is the most prevalent primary malignant brain tumor and remains uniformly fatal despite aggressive therapies including surgery, radiation, and chemotherapy ^1, 2^. With limited treatment options, the success of immunotherapies in other advanced cancers, including melanoma and non-small cell lung cancer, has inspired investigation of immune based therapies in GBM ^3–6^. However, early clinical trials of immune checkpoint therapies in GBM have demonstrated limited response, if any, and despite some evidence of immune cell accumulation, GBM growth persists ^7, 8^. One explanation for these failures could be the potent immunosuppressive factors present in GBM, including the high tumor content of myeloid-derived suppressor cell (MDSC) ^9–12^. MDSCs are a heterogeneous population of bone marrow-derived cells consisting of monocytic (M-MDSC) and granulocytic (G-MDSC) subsets that accumulate in the tumor, spleen, and peripheral blood of GBM patients, where they exert immune suppression by dampening the function of natural killer (NK) cells and cytotoxic T lymphocytes (CTLs) ^13–18^.

Recent work from our laboratory and others identified an increase in circulating M-MDSCs in the peripheral blood of GBM patients compared to benign and grade I/II glioma patients^9, 19^. However, this difference was not observed for other immunosuppressive cell populations, such as macrophages or T-regulatory cells, which were not different between patients of different glioma grades. In addition, MDSCs in the peripheral circulation and infiltrating in the GBM microenvironment correlated with poor prognosis^9, 19^. Based on these observations in GBM and other cancers, attempts to target MDSCs using multiple approaches, including low-dose chemotherapy in a recent GBM trial are in clinical evaluation^20^. Notably, these approaches use non-specific strategies that attenuate MDSCs, as opposed to targeted approaches that are MDSC-specific and may have a higher therapeutic utility.

In seeking to develop MDSC targeted therapies to reduce immune suppression, we focused our attention on macrophage migration inhibitory factor (MIF). Targeting MIF is of interest due to our previous work where we observed that MIF derived from GBM cells, specifically therapeutically resistant cancer stem cells (CSCs), was necessary for MDSC survival and function^21^. Moreover, reducing MIF levels in GBM cells did not alter their proliferation, but when transplanted into an immune competent orthotopic model, resulted in increased host survival and an increase in the number of CD8 T cells in the tumor microenvironment. MIF has also been shown by other groups to enhance the immune suppressive capacity of myeloid cells^22^; for instance, MIF downregulation was demonstrated to aid in the resistance of anti-VEGF therapies^23^. In seeking to understand exactly how MIF effects the immune response in GBM one must consider that it has been shown to be highly context specific, exerting both inflammatory and anti-inflammatory effects depending on the disease and tissue^21, 24−28^. MIF signals through a variety of receptors, including via its cognate receptor CD74, and by non-cognate interactions with CXCR2, CXCR4, CXCR7. CD74 is the cell surface form of the Class II invariant chain, but is expressed independently of Class II to mediate MIF signal transduction^29–31^. MIF binding to CD74 leads to the recruitment of CD44 as a signaling co-receptor, leading to downstream pERK signaling. By contrast, MIF signaling through CXCR2 primarily activates PI3K/AKT pathways^32^. The pharmacologic targeting of MIF has also been of great interest in a variety of inflammatory conditions including multiple sclerosis, systemic lupus erythrematosus, rheumatoid arthritis, inflammatory bowel disease, and other inflammatory disorders^22, 33–40^. Additionally, clinically approved MIF inhibitors have been developed that could potentially be repurposed for GBM^33^. To gain a more mechanistic understanding into the MIF signaling axis in MDSCs for potential targeting GBM, we examined the expression and function of MIF receptors in MDSCs derived from mouse and human GBM models.

## Methods

### Co-culture assay

Co-culture induction of MDSCs was adapted from previously described work in melanoma^*28*^. At day zero bone marrow (BM) was freshly isolated from the tibias and femurs of male 000664-C57BL/6J. To obtain BM derived MDSCs, the freshly isolated BM was incubated for 3 days in a medium consisting of 50% conditioned medium from a 24hr GL261 (glioma) cell line culture and fresh RPMI medium with 10% FBS. Additionally, this medium was supplemented with GM-CSF (40 ng/mL, Biolegend Catalog # 575906), and IL-13 (80 ng/mL, Biolegend Catalog # 576306), which have been shown to increase MDSC expansion and activity. BM was cultured in this medium in 6 well plates at a density of 2,000,000 cells per well as previously described and utilized for analysis on day 3 post initiation^28^.

### Flow cytometry of co-culture

At day 3 of the co-culture cells were extracted from the wells using gentle washing with RPMI medium, blocked in FcReceptor block (Miltenyi Biotec 130-092-575) and then stained live on ice. Samples were then fixed using eBioscience fixation buffer before analysis. Gating for MDSCs was performed using FlowJO V10, and M-MDSCs were identified by (Singlets/Live/CD45+/CD11b+/CD68−/IAIE−/Ly6G−/LyC+) and G-MDSCs by (Singlets/Live/CD45+/CD11b+/CD68−/IAIE−/Ly6C−/Ly6G+). Antibodies were obtained from Biolegend (San Diego, CA) for analysis of mouse immune profile Fluorophore-conjugated anti-Ly6C (Clone HK1.4, Catalog # 128024), anti-Ly6G (Clone A8, Catalog # 127618), anti-CD11b (Clone M1/70, Catalog # 101212), anti-CD68 (Clone FA-11, Catalog # 137024), anti-I-A/I-E (Clone M5/114.15.2, Catalog # 107606), anti-CD11c (Clone N418, Catalog # 117330), anti-Ki-67 (Clone 16A8, Catalog # 652404), anti-CD45 (Clone 30-F11, Catalog # 103132), anti-CD74 (Clone IN-1 Catalog # 740385), anti-P2Ry12 (Clone S16007D, Catalog # 848004), anti-CXCR2 (Clone SA044G4, Catalog # 149313), anti-CXCR4 (Clone L276F12, Catalog # 146506), anti-CXCR7 (Clone 8F11-M16, Catalog # 331115), anti-CD44 (Clone IM7, Catalog # 103039). Compensation was performed using AbC Total Antibody Compensation Bead Kit (Catalog # A10497).

### Flow Cytometry Patient Tumor Samples

Flow cytometry data was utilized from *D. Peereboom T. Alban et al., JCI insight 2019 ^20^*. Tumor tissue was received from recurrent GBM patients undergoing treatment in clinical trial NCT02669173. Tissue was digested in collagenase IV (STEMCELL Technologies) for 1 hour at 37 degrees Celsius and then mechanical dissociated via 40-uM filter. Dissociated tumors were then washed in RPMI medium before being viably frozen for flow cytometry analysis. MDSC panel consisted of CD11b (Catalog # CD11b29), HLA-DR (Catalog # 559866), CD14 (Catalog # 560180), CD15 (Catalog # 555400), CD33 (Catalog # 555450), CXCR2 (Catalog # 551126), CD74 (Catalog # 555538 with Lightning-Link PE-Cy7 Catalog # 762-9902). Staining and analysis were performed using standard protocols previously described, with MDSCs marked by CD11b+, CD33+, and HLA-DR–/lo and then further subdivided into granulocytic MDSCs (CD15+) and monocytic MDSCs (CD14+)^9, 20, 41^. After gating for MDSC populations the MFI of CXCR2 and CD74 was analyzed using FlowJo V10 for each sample.

### T cell suppression assay

At day 3 post MDSC co-culture, T cell suppression was assessed. Splenocytes were freshly isolated from male 000664-C57BL/6J mice using sterile techniques. Post isolation the red blood cells were lysed using RBC lysis buffer (Biolegend Catalog # 420301) before being magnetically sorted using the (Pan T cell isolation kit Catalog # 130-095-130, Miltenyi Biotec). Isolated T cells were then stained using CFSE Cell Division Tracker Kit (Biolegend Catalog # 23801). CFSE stained T cells were then collected and distributed into round bottom 96 well plates at 100,000 cells per well in IL-2(30 IU) as unstimulated control. Stimulated controls additionally contained CD3/CD28 mAb-coated beads (ThermoFisher Scientific) at a ratio of 3:1. T-cell activation was measured by flow cytometry with the controls consisting of CFSE labeled T cells alone and CFSE labeled T cells with beads. Co-culture derived MDSCs, isolated by magnetic sorting (MACS Miltenyi MDSC isolation kit Catalog # 130-094-538), were seeded with T cells at a concentration of 1:2 (1 MDSC for every 2 T cells).

### Quantitative PCR

Quantitative PCR was performed for MDSC markers and immune suppressive genes Arg1 (Forward: AAGAATGGAAGAGTCAGTGTGG, Reverse: GGGAGTGTTGATGTCAGTGTG), iNOS (Forward: TGTGCTTTGATGGAGATGAGG, Reverse: CAAAGTTGTCTCTGAGGTCTGG), Ly6G (Forward:TTGTATTGGGGTCCCACCTG, Reverse: CCAGAGCAACGCAAAATCCA), CXCR2 (Forward: TCTTCCAGTTCAACCAGCC, Reverse: ATCCACCTTGAATTCTCCCATC), CD74 (Forward: ATGGCGTGAACTGGAAGATC, Reverse: CAGGGATGTGGCTGACTTC), MCP-1 (Forward: GTCCCTGTCATGCTTCTGG, Reverse: GCTCTCCAGCCTACTCATTG).

RNA was isolated using Qiagen RNeasy Mini Kit and cDNA was generated using aScript cDNA SuperMix (Quantabio). After cDNA generation qPCR was performed using the Fast SYBR™ Green Master Mix (ThermoFisher Scientific).

### GBM-seq database mining

Darmanis, S et al., Cell Reports 2017data was utilized in this analysis where normalized count data was acquired from GBMseq.org^42^. Subsequently, CD74 and other MIF receptor expression levels were graphed for the myeloid populations and other immune populations as characterized by Darmanis, S et al. in their supplemental data. All populations’ names were kept the same as previously published and identified.

### MIF inhibitor screen

The co-culture system was utilized to screen inhibitors of MIF and MIF/CD74 interaction by dosing inhibitors at day zero when the co-culture was initiated and then reading out % MDSCS of live cells by flow cytometry. The same gating strategy as in the co-culture methods section was used to determine if the MDSC population was shifting. Screens were performed in biological replicates of 3 on two separate experiments for a total of 6 biological replicates. The studied MIF inhibitors were anti-MIF mAb (IIID.9), 4-IPP (Tocris Catalog # 3429)^43^, Ibudilast (gift of Medicinova)^44–46^, ISO-1 (Tocris Catalog # 4288)^43^, MIF098^47–49^, AV1013(gift of Medicinova)^46^, and the PDE4 inhibitor was Rolipram (Tocris Catalog # 0905).

### In vivo syngeneic glioma model

Ibudilast treatment was assessed in two cohorts using the syngeneic mouse model of glioma GL261 acquired from the NCI. 6-week-old aged-matched male 000664-C57BL/6J mice were anesthetized using isoflurane and then intracranially injected in the left cerebral hemisphere with 20,000 GL261 cells in 5 μl of RPMI medium using a stereotactic frame. This model has been established in the laboratory with neurological symptoms as an indicating endpoint and a median survival time of approximately 20 days^21^. Flow cytometry was performed on mechanically dissociated tumors isolated from the left hemisphere from sacrificed animals at day 18 post implantation, and a terminal cardiac bleed was analyzed for MDSC and T cell levels using the myeloid panel: live/deadUV, CD45, CD11b, CD11C, IA/E, CD74, Ly6G, Ly6C, CD68, and the lymphoid panel: live/deadUV, CD45, CD3, CD4, CD8, PD1, NK1.1, CD107a. Antibodies were obtained from Biolegend (San Diego, CA) for analysis of mouse immune profile Fluorophore-conjugated anti-Ly6C (Clone HK1.4, Catalog # 128024), anti-Ly6G (Clone A8, Catalog # 127618), anti-CD11b (Clone M1/70, Catalog # 101212), anti-CD68 (Clone FA-11, Catalog # 137024), anti-I-A/I-E (Clone M5/114.15.2, Catalog # 107606), anti-CD11c (Clone N418, Catalog # 117330), anti-CD3 (Clone 145-2C11, Catalog # 100330), anti-CD4 (Clone GK1.5, Catalog # 100422), anti-CD8 (Clone 53-6.7, Catalog # 100712), anti-NK1.1 (Clone PK136, Catalog # 108741), anti-CD45 (Clone 30-F11, Catalog # 103132). An initial study included 10 vehicle and 10 Ibudilast treated animals, but at day 18, the 2 vehicle treated animals demonstrated neurological symptoms and were euthanized prior to analysis time-point. Additionally, tumor could not be identified visually at day 18 in 3 ibudilast treated mice and 2 vehicle treated mice, so their matched non-tumor bearing tissue was not included in analysis.

### Nanostring analysis

RNA was isolated using RNeasy mini kit (Qiagen) and then the nCounter® Mouse Myeloid Innate Immunity Panel v2 was used to analyze the RNA expression of tumors isolated from 6 endpoint vehicle tumors and 6 endpoint Ibudilast treated animals.

### Immunohistochemically analysis

At endpoint, vehicle and ibudilast treated animals were perfused with 4%PFA before removing the brain and fixing in PFA overnight at 4 °C. Post Fixed brains were cryopreserved in sucrose and embedded in O.C.T compound (Fisher Healthcare) to make frozen sections (10μm thick). Endogenous peroxide activity was quenched by 3% H2O2 incubation and blocked in 5% normal goat serum/0.2%Triton in PBS for 30 minutes before primary antibodies were added. Phospho-Histone3 (1:500, catalog # 06-570, MillopreSigma) and Ki67 (1:1,000, catalog # ab15580, Abcam) antibodies were allowed to bind overnight at 4 °C. After rinsing with 1xPBS, biotinylated secondary antibodies (1:500, Invitrogen) were added and incubated at RT for 1hr. Signal was amplified using avidin-biotin complex staining (30mins) before DAB substrate was used to visualize the signal (Vector Laboratories). Hematoxylin was used for counterstain. After washing in PBS, the slides were dehydrated through alcohol series and mounted with Permount (Fish Chemical).

### Statistical analysis

Graph-Pad Prism was utilized for statistical analysis of survival curves for log-rank tests and also for T-tests throughout the manuscript. *<0.05, ** <0.01, ***<0.001. Nanostring statistics were performed within nSolver software supplied by Nanostring and the advanced analyzer V 4.0.

## Results

### Development of MDSC co-culture to study the MIF signaling axis

While MDSCs have been linked to GBM prognosis and progression, technical hurdles including the inability for their long-term expansion have been a challenge for mechanistic insight and functional assessment^9, 19, 50^. Previously our group identified MIF secreted by GBM CSCs and driving MDSCs, however the mechanism by which MIF increased MDSC function remains unclear^21, 22^. Initially we sought to determine if the survival extension we previously observed with MIF knockdown GBM cells was solely due to an immunologic event. We performed the same studies in immune compromised NSG mice and found that there was no survival benefit when the adaptive immune response was absent (**Supplemental Figure 1A-B**). Furthermore, when MIF was depleted using an established neutralizing anti-MIF antibody 5-days post tumor implantation there was no survival benefit (**Supplemental Figure 1C**). These findings confirm our previous observations that MIF likely acts on the immune system, as opposed to acting on GBM cells in an autocrine manner. To further understand how GBM-derived MDSCs function, we adapted a co-culture system previously developed in a melanoma model (**Figure 1A**) ^28^. The co-culture utilizes freshly-isolated bone marrow combined culture in 50% conditioned media from a 24-hour culture of the mouse glioma cell line GL261 and supplemented with GM-CSF and IL-13 to generate M-MDSCs and G-MDSCs over a 3-day period. Day 3 was chosen for MDSC generation assays based on a flow cytometry longitudinal study of the culture showing a steep decline in viable CD45+ cells after day 4 (**Supplemental Figure 1D**). At day 3 of co-culture, the numbers of M- and G-MDSCs were determined by flow cytometry analysis where M-MDSCs were gated by Singlets/Live/CD45^+^/CD11b^+^/CD68^−^/IAIE^−^/Ly6G^−^/Ly6C^+^ and G-MDSCs by Singlets/Live/CD45^+^/CD11b^+^/CD68−/IAIE^−^/Ly6C^−^/Ly6G^+^. Furthermore, co-culture generated MDSC function was determined by T cell suppression assay. In this assay, CFSE labeled T cells which were activated by CD3/CD28 mAb coated beads, were suppressed by MDSCs at a ratio of 1 MDSC to 2 T cells (**Figure 1B**). Furthermore, FACs sorted M-MDSCs and G-MDSCs were analyzed by QPCR for Arginase-1, iNOS, and Ly6G to validate the subsets, and G-MDSCs were observed to have increased Ly6G and iNOS, while M-MDSCs highly expressed Arginase-1 (**Figure 1C**). These data validate a model system for generating functional GBM-educated MDSCs as a platform for functional assessment and inhibitor studies.

**Figure 1.**
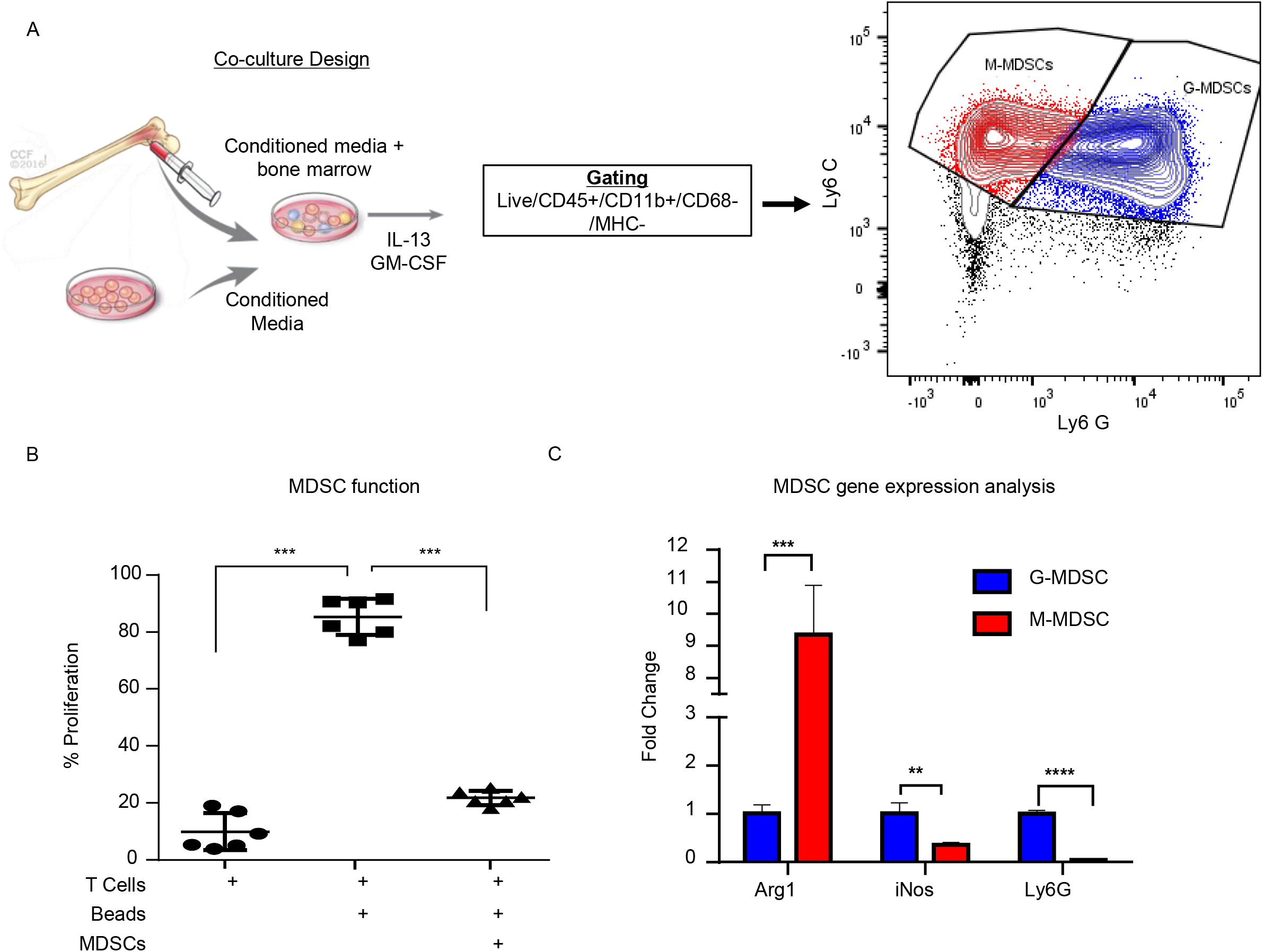
Glioma educated MDSCS can be generated in vitro. MDSCs are induced using freshly isolated bone marrow cultured with 50:50 mix of fresh media and conditioned media from a 24-hour culture of GL261 cells with the addition of IL-13 and GM-CSF over 3 days (**A**). M-MDSCs were gated by Live/CD45^+^/CD11b^+^/CD68^−^/MHC^−^/Ly6C^+^/Ly6G^−^ while G-MDSCs were gated by Live/CD45^+^/CD11b^+^/CD68^−^/MHC^−^/Ly6C^+^/Ly6G^+^. Co-cultured MDSCs from n=6 mice were generated over 3 days and then isolated by magnetic bead sorting and subsequently used for T cell suppression assay where the controls were T cells alone unstimulated without CD3/CD28 activation beads and T cells with CD3/CD28 activation beads (**B**). FACs sorting of M-MDSCs and G-MDSCs from 3 day old co-cultures of n=3 mice was used to isolate RNA and perform qPCR for Arginase (Arg1), Nitric oxide synthase (iNOS), and Ly6G (**C**). Two-Tailed T-Test was performed for comparisons in panel B, C *<0.05, ** <0.01, ***<0.001.

### In vivo and in vitro analysis demonstrate M-MDSCs with surface expression of the MIF receptor CD74

In order to determine the MDSC subset driving immune suppression GBM, we used a syngeneic model of glioma GL261, which was intracranially implanted to generate syngeneic tumors. At day 18 post implantation the tumor bearing (left) and non-tumor bearing (right) hemispheres were removed and analyzed by flow cytometry for MDSC subpopulations using the same gating strategy as in the co-culture system with the addition of pP2RY12 to exclude microglia. Analysis identified higher levels of M-MDSCs in the tumor bearing and non-tumor bearing hemispheres of the brain compared to G-MDSCs (**Figure 2A**). In order to determine the MIF receptor profiles, flow cytometry of the MIF receptors CD74, CXCR2, CXCR4, and CXCR7 was performed 3-days post co-culture initiation (**Figure 2B, C**). The percent positive for each receptor was analyzed by flow cytometry, which identified M-MDSC as having high expression of CD74 and its co-receptor CD44, while G-MDSCS primarily expressed CXCR2 (**Figure 2B, C**). FACs sorted M-MDSCs and G-MDSCS from co-cultures confirmed these findings, showing CXCR2 expression in G-MDSCs, and CD74 with the downstream effector MCP-1 as being highly expressed, suggesting activation through MIF/CD74 signaling axis (**Figure 2D**)^51^. Furthermore, the analysis of M-MDSCs by flow cytometry showed high levels of CD74 expression compared to G-MDSCs (**Figure 2E**), and when quantified significantly higher than in G-MDSCs (**Figure 2F**). Interestingly, when MDSCs were permeabilized and stained for intracellular CD74 there was no difference between G- and M-MDSCs in the intracellular amounts of CD74 (**Figure 2G**). In vivo analysis of M-MDSCs in the tumor microenvironment using the syngeneic glioma model further supports these findings by showing the mean fluorescence intensity (MFI) of CD74 as higher on M-MDSCs compared to G-MDSCs or microglia of the tumor bearing hemisphere (**Figure 2H**). Taken together, these data demonstrate differential MIF receptor expression in MDSC subsets in mouse models.

**Figure 2.**
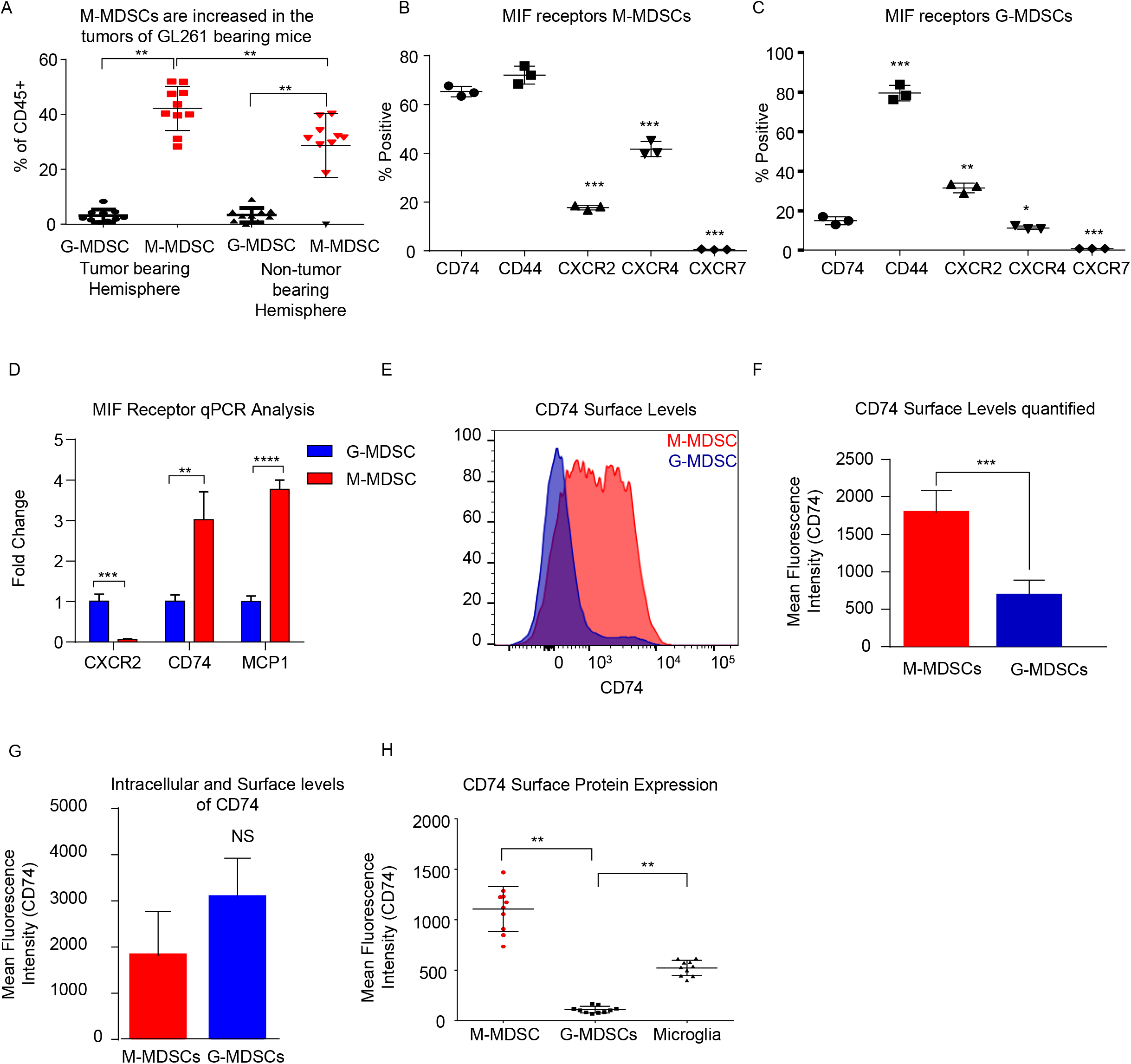
Murine M-MDSCs express the MIF receptor CD74. n=10 mice were intracranially injected with the syngeneic mouse glioma cell line GL261 at day 0 and then at Day 18 post injection the tumor bearing and non-tumor bearing hemispheres were resected, dissociated and analyzed by flow cytometry (**A**). M-MDSCs Live/CD45^+^/CD11b^+^/CD68^−^/P2Ry12−/MHC^−^/Ly6C^+^/Ly6G^−^, and G-MDSCs Live/CD45^+^/CD11b^+^/CD68^−^/P2Ry12-/MHC^−^/Ly6C^+^/Ly6G^+^. n=3 mice were used for co-culture induction of MDSCs and at day 3 M-MDSCs and G-MDSCs were analyzed for the MIF receptors CD74, CD44, CXCR2 CXCR4, and CXCR7 by flow cytometry and gated for the % positive in each group (**B, C**). FACs sorting of G-MDSCs and M-MDSCs from co-cultures of n=3 mice were performed and then RNA isolated for qPCR analysis of the expression of MIF receptors (CXCR2, and CD74) as well as MCP-1, the CD74 downstream activation product (**D**). CD74 expression was assessed by flow cytometry using flow cytometry staining of co-cultures where the histogram demonstrates the expression level of CD74 on M-MDSCs compared to G-MDSCs (**E**). Quantification of n=3 co-culture derived M-MDSCs and G-MDSCs CD74 Mean Fluorescence intensity shows higher levels of CD74 on M-MDSCs (**F**). Intracellular staining post permiablization of the same cohort of M-MDSCs and G-MDSs from panel (**F**) shows that CD74 levels were not significantly different when staining internally (**G**). In vivo, the tumor bearing mice that were evaluated for MDSC levels in panel (A) were also evaluated for CD74 expression on the surface of M-MDSCS, G-MDSCS, and Microglia (**H**). Two-Tailed T-Test was performed for comparisons in panel A,D, F, G, H *<0.05, ** <0.01, ***<0.001.

### GBM patient derived specimens show the MIF receptor CD74 expressed on MDSCs and associate with poor prognosis

To determine if the findings in the mouse glioma model are recapitulated in the tumor microenvironment of human GBM patients, we utilized bioinformatics analysis of previously published single-cell sequencing datasets and flow cytometry analysis of GBM tumor specimens. The GBMseq dataset provides single cell sequencing of 3,589 cells from a cohort of 4 GBM patients annotated for their population names ^42^. Utilizing this dataset, we isolated the log2 counts for the myeloid populations identified and looked at the MIF receptor expression of CXCR2, CXCR4, CXCR7, CD74, and CD44 (**Figure 3A**)^31^. Statistical analysis revealed that CD74 was most highly expressed in the myeloid populations. Furthermore, using the annotated populations, the level of CD74 expression was compared across all populations in the GBMseq dataset, which revealed highest levels on the myeloid cells (**Figure 3B**). To validate these findings, a separate cohort of 8 GBM tumors were analyzed by flow cytometry using a human panel previously validated, where M-MDSCs were identified by the following gating strategy singlets/live/HLA-DR^−^/CD33^+^/CD11b^+^/CD14^+^/CD15^−^ and G-MDSCs by singlets/live/HLA-DR^−^/CD33^+^/CD11b^+^/CD14^−^/CD15^+^. The expression of CD74 and CXCR2 were analyzed on reach subpopulation by MFI, where CD74 was shown to be more highly expressed on M-MDSCs, while CXCR2 was more highly expressed on G-MDSCs (**Figure 3C**). Based on these findings, we tested the hypothesis that MIF and CD74 are signaling together and driving GBM immune suppression. We analyzed the cancer genome atlas (TCGA) GBMLGG database for survival and MIF expression and CD74 expression and the combination (**Figure 3E-G**). These data demonstrate that MIF and CD74 expression individually predict a poor prognosis, but when combined into MIF and CD74 double high as defined by greater than median expression of MIF and CD74, then the prognosis becomes poorer as demonstrated by hazard ratios MIF alone HR: 1.51, CD74 alone HR: 1.69, MIF/CD74 HR:2.45 (**Figure 3G**). These data demonstrate that human GBM specimens’ express high levels of CD74 in M-MDSCs in the tumor microenvironment that are linked to poor patient prognosis.

**Figure 3.**
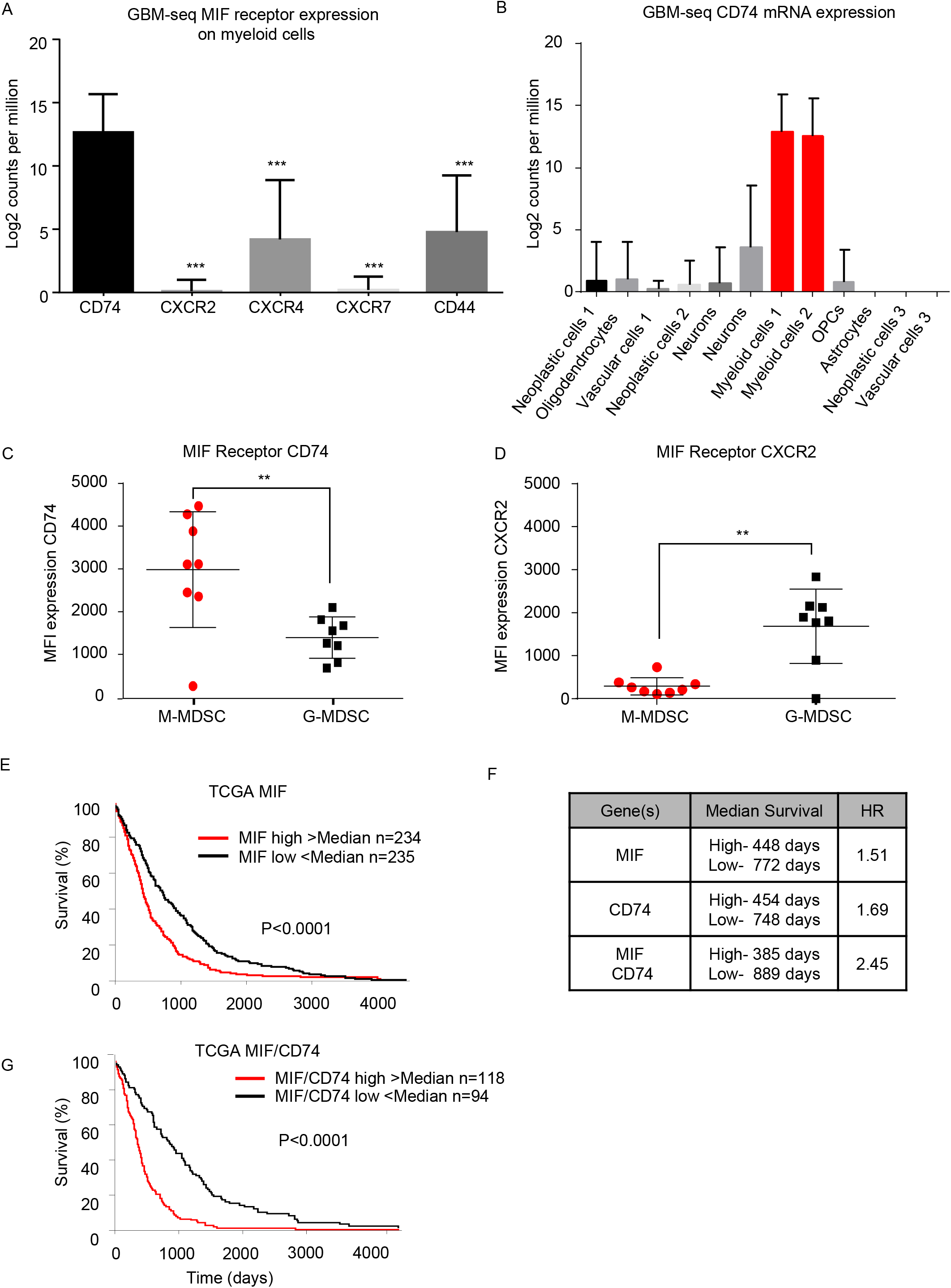
Human derived M-MDSCs express the MIF receptor CD74. Data mining of the GBM-seq database from Darmanis, S et al., Cell Reports 2017 was used to analyze the myeloid cell expression of the MIF receptors CD74, CXCR2, CXCR4, CXCR7 and CD44 showing that CD74 expressed by the myeloid populations in GBM tumor single cell sequencing (**A**). Further analysis was performed separating the single cell populations into the previously published cell identities (**B**). Using a previously published cohort of GBM patient tumors^20^ n=8 GBM patients the MIF receptors CD74 and CXCR2 were assessed on M-MDSCs and G-MDSCs (M-MDSCs: CD11b^+^/ HLA-DR^−^/ CD33^+^/ CD14^+^/CD15^−^, G-MDSCs: CD11b^+^/ HLA-DR^−^/CD33^+^/ CD14^−^/CD15^+^) (**C,D**). TCGA data analysis of GBMLGG cohort identified MIF expression and CD74 expression levels correlating with survival with a similar hazard ratio (HR) (**E, F**). When a signature for both MIF and CD74 is created where samples that were above the median for both MIF and CD74 expression compared to those below the median for both MIF and CD74 further separates survival (**F, G**). Two-Tailed T-Test was performed for comparisons in panel A, C, D *<0.05, ** <0.01, ***<0.001. Survival curve analysis was performed in GraphPad Prism using Log-rank (Mantel-Cox) test for p value and hazard ratio log rank was computed on the same data using GraphPad Prism.

### MIF inhibitor screening identified the MIF/CD74 interaction inhibitor Ibudilast

In order to identify a potential targeted therapy that acts on the MIF/CD74 signaling axis and neutralizes M-MDSCs, we utilized the in vitro co-culture system to generate glioma educated MDSCS in the presence of different small molecule MIF inhibitors. In this system the generation of M-MDSCs was monitored at day 3 post co-culture in the presence of various MIF inhibitors at 200 μM, a concentration achieved reached in circulation with Ibudilast, a primary drug of interest due to its known toxicity profile and ability to penetrate the blood brain barrier (**Figure 4A**)^45, 52^. Other MIF inhibitors previously identified as either MIF tautomerase inhibitors (4-IPP, ISO-1, AV1013, MIF098), or MIF/CD74 interaction inhibitors (Ibudilast, MIF098), were compared to Ibudilast at similar 200 μM concentrations to determine the specificity of ibudilast in reducing M-MDSCs^43, 46, 49^. The MIF/CD74 interaction inhibitor Ibudilast demonstrated an effective reduction in M-MDSC generation (**Figure 4A**). This reduction in M-MDSCs was not a result of a major change in cell viability as assessed by live/dead staining. Additionally, the MIF inhibitor 4-IPP, which does disrupt the interaction of MIF with CD74 showed no efficacy (**Figure 4A**)^43^. While ibudilast has been shown to inhibit the interaction of MIF and CD74, there are also reports that it is a phosphodiesterase inhibitor^53, 54^. To assess specificity, we compared Ibudilast at 100 μM and 200 μM to Rolipram, which is a known specific and potent phosphodiesterase inhibitor at the same concentrations (**Figure 4B**)^55^. Rolipram was unable to alter the generation of M-MDSCs and thus the reduction of M-MDSCs is likely not due to these off-target effects of ibudilast. The reduction in M-MDSC generation was not a result of a major change in cell viability as assessed by live dead staining. Also, to determine if MDSCs could be killed by Ibudilast an IC-50 was attempted using FACs sorted M-MDSCS and G-MDSCs increasing doses of Ibudilast were added to cultures for 24 hours before being analyzed. No change in viability of M- or G-MDSCs was detected (**Figure 4D**). The function of MDSCs generated in co-culture with Ibudilast was analyzed using the T cell suppression assay, and identified as a reduction in the ability of MDSCs to suppress T cell proliferation (**Figure 4E**). Additionally, untreated M-MDSCs and G-MDSCs were isolated by FACs sorting and then treated for 24 hrs with Ibudilast before western blot analysis for pERK, a proximal downstream target of MIF/CD74 signaling^51^. This revealed a specific reduction of pERK signaling compared to total ERK expression in M-MDSCs and not in G-MDSCs, showing specific MIF/CD74 inhibition by ibudilast in M-MDSCs (**Figure 4F**). These data demonstrate that M-MDSC generation and function and be disrupted by pharmacologic a MIF/CD74 inhibition.

**Figure 4.**
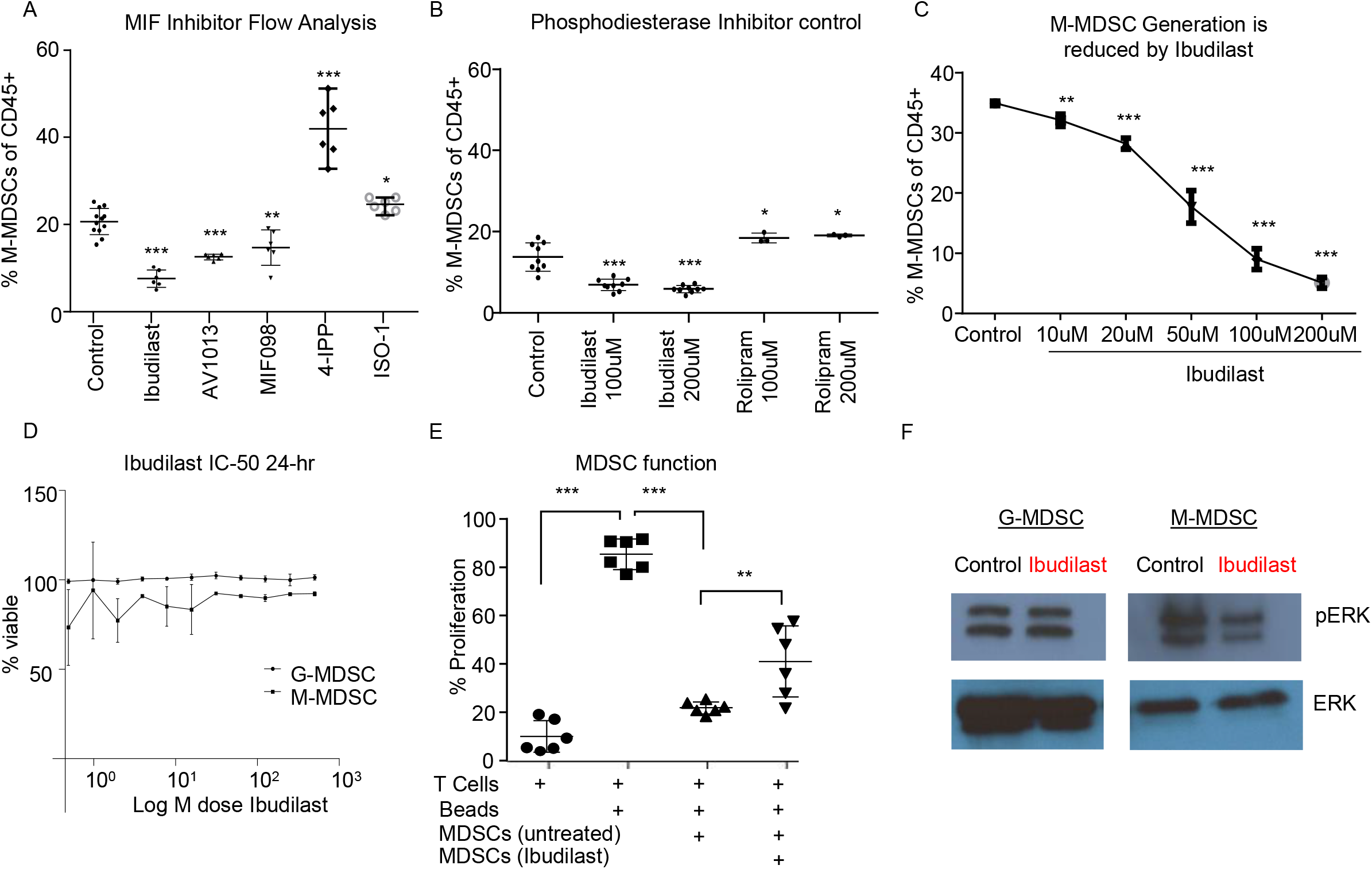
Ibudilast inhibits the MIF disrupting M-MDSC generation in vitro. Utilizing the co-culture system described in **Figure 1** MIF inhibitors were assessed for their ability to inhibit MDSC generation (**A**). Inhibitors were added at 200uM at day 0 during the co-culture initiation and then assessed at day 3 for the % of M-MDSCs of CD45+ cells (**A**) n=6 mice from n=2 separate experiments. As a control for Ibudilast off target effects on phosphodiesterase Ibudilast was directly compared to Rolipram at 100uM and 200uM doses n=6 control and Ibudilast treated co-cultures from n=6 mice and n=3 Rolipram treated co-cultures (**B**). n=3 mice per co-culture were used and ibudilast evaluated at 10uM, 20uM, 50uM, 100uM and 200uM and then assessed by flow cytometry at day 3 (**C**). To determine if ibudilast was killing the M-MDSCs or G-MDSCs we isolated M-MDSCs and G-MDSCs from untreated co-cultures at day 3 from n=3 mice by FACs sorting and then treated them for 24hours with Ibudilast as an IC50 using celltiterglo as a readout for viability (**D**). The function of MDSCs treated with ibudilast was assessed by generating MDSCs in the presence of ibudilast and then magnetically sorting for MDSCs comparing untreated and Ibudilast treated MDSCs (**E**). To assess the disruption of the MIF/CD74 signaling mechanism M-MDSCs and G-MDSCs were FACs sorted from day 3 co-cultures and then subsequently 50ng/ml MIF was added to each well containing 500,000 cells and then treated with Ibudilast at 200uM for 24 hours prior to lysing the cells and performing western blot analysis for pERK and total ERK (**F**). Two-Tailed T-Test was performed for comparisons in panel A, B, D, E *<0.05, ** <0.01, ***<0.001.

### Ibudilast treatment reduced MIF/CD74 signaling in a syngeneic model

To determine the in vivo effects of Ibudilast treatment, a cohort of tumor bearing animals were treated 5 days post tumor implantation (at 50 mg/kg 2x weekly based on previous experiments and the known effect dose effect of Ibudilast in a murine model ^45^). Daily dosing has been demonstrated in rodents to increase CYP enzymes and degrade ibudilast, reducing the bioavailability^45^, and thus high doses of bi-weekly ibudilast was chosen for this treatment. Animals were analyzed at endpoint and tumors were dissected from the brain for RNA analyses by Nanostring Ncounter myeloid panel. Initial analysis by principal component analysis revealed that vehicle tumors and ibudilast tumors separate and the separation is driven by the vectors of MIF, CD74, PTGS2, Arg1, CXCR2 (**Figure 5A**). A volcano plot comparing the significantly differentially expressed genes between vehicle and ibudilast treated tumors showed significant change in immune genes upon treatment (**Figure 5B**). Pathway analysis between vehicle and Ibudilast treated tumors showed reduced antigen presentation, which coincides with reduced CD74 and MHC expression following the hypothesis that Ibudilast is targeting CD74 in vivo as well as in vitro (**Figure 5C**). Pathway analysis also demonstrated increased Lymphocyte activation upon treatment showing possibly increased immune response (**Figure 5C**) and CD74 expression was reduced upon treatment as expected (**Figure 5D**). Furthermore, analysis of Nanostring data also revealed a predicted reduction of MEK2 expression, which is downstream of MIF/CD74 signaling (**Figure 5E**) and consistent with the western blot findings of reduced pERK signaling upon Ibudilast treatment. Flow cytometry analysis of tumor, non-tumor, and blood from this cohort at day 18 post injection tumors, 14 days of Ibudilast treatment, identified an increase in CD8 T cells specific to the tumor, while other immune cell populations were unchanged (**Figure 5F**, **Supplemental Figures 2,3**). Additionally, immunohistochemistry staining identified a reduction of proliferation in Ibudilast treated tumors via reduced p-Histone3 and ki-67 staining (**Supplemental Figure 4**). Importantly, we saw no changes in other T cell or myeloid cell populations, including the overall number of CD45+ cells (**Supplemental Figures 2,3**). Taken together, these data reveal that CD74/MIF inhibition via Ibudilast can reduce MDSCs in vivo and increase immune activation in the tumor microenvironment.

**Figure 5.**
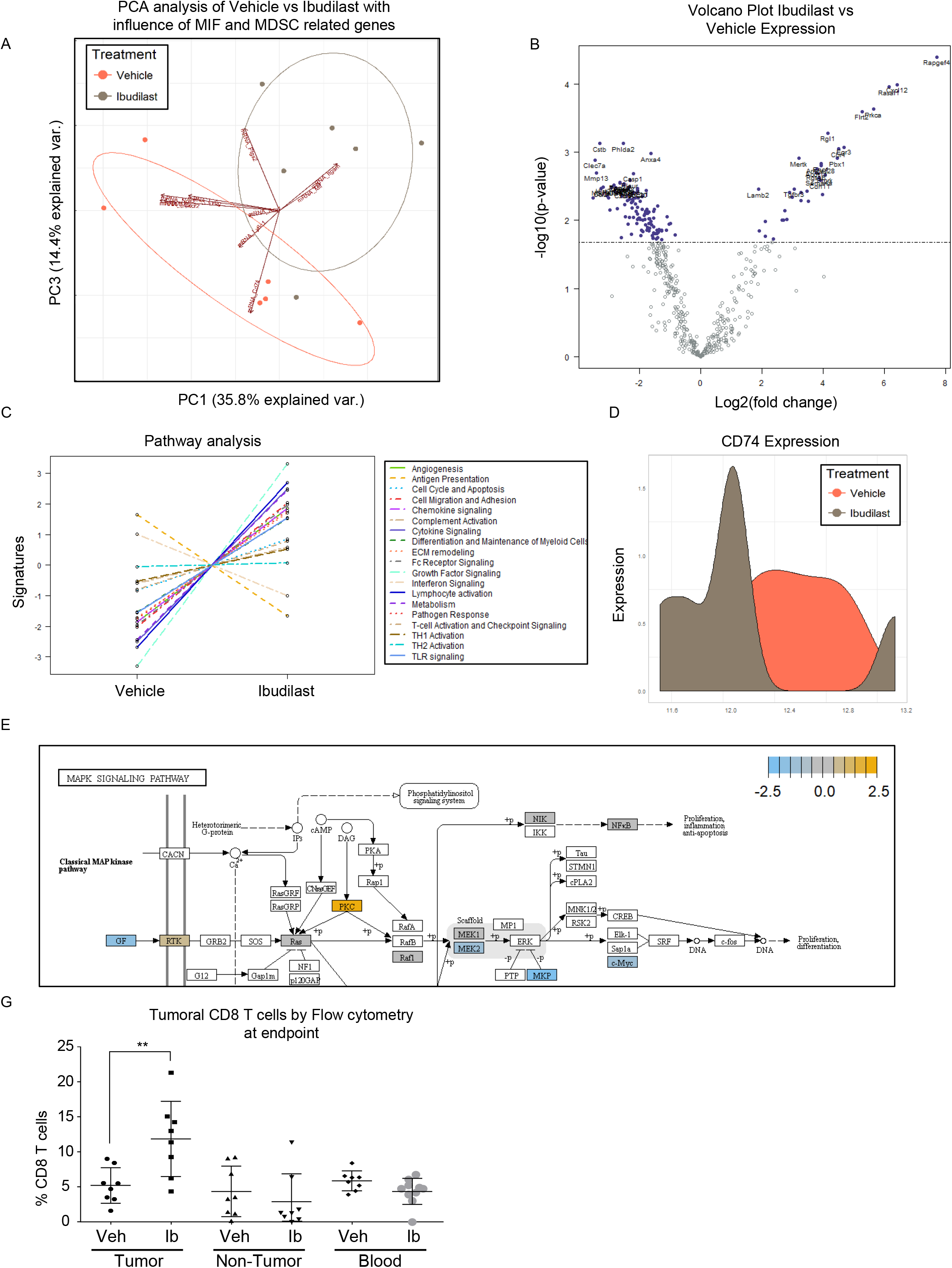
Ibudilast inhibits the MIF disrupting M-MDSC generation in vitro. n=6 vehicle and n=6 Ibudilast treated mice (50mg/kg 2x weekly starting day 5 post tumor implantation) were sacrificed at endpoint and tumors were dissected from the brain for RNA isolation. RNA from isolated tumors of vehicle and ibudilast treated mice was analyzed via Nanostring murine myeloid panel and PCA was performed showing separation of ibudilast vs vehicle (**A**). Volcano plot comparing log2fold change in genes between Ibudilast and vehicle demonstrates significant changes in the myeloid populations between vehicle and ibudilast treated tumors (**B**). Pathway analysis of Ibudilast treated tumors shows increased activation of many immune pathways including lymphocyte activation while there is a reduction in antigen presentation (**C**). Summary of CD74 expression in histogram format comparing all Vehicle and all Ibudilast treated samples (**D**). Pathway analysis of Nanostring data identifies the MAPK signaling pathway in Ibudilast treated tumors with a reduction in MEK2 (**E**).. A cohort of n=8 vehicle and n=8 Ibudilast treated mice were sacrificed at day 18 post injection and tumor, non-tumor tissue, and blood were analyzed by flow cytometry for immune populations where CD8 T cells were shown to be significantly increased in the tumors of Ibudilast treated mice (**F**). Two-Tailed T-Test was performed for comparisons in panel F *<0.05, ** <0.01, ***<0.001. All other statistics were performed in Nanostring Nsolver software including the PCA and volcano plot differential gene expression and pathway analysis.

## Discussion

While multiple groups including our own have identified MDSCs as being increased in GBM and other cancers^9, 11, 12, 19, 21^, our understanding of the factors driving these cells has been lacking and strategies to target these cells has not matured. Here we focused our efforts on MIF as a driver of MDSCs based on our previous work showing that MIF depletion could reduce MDSC function^21^. Additionally, multiple groups have indicated a link between MIF and MDSCs^22, 28, 56^. Here we found that the receptor CD74 may play a greater role in GBM MDSC biology because the subset of MDSCs primarily found in the tumor microenvironment were M-MDSCs, which primarily express CD74 as a MIF receptor. This is in contrast to metastatic breast cancer models that show G-MDSCs infiltrating tumors and driving metastasis^57, 58^; where in those cases we would hypothesize that CXCR2 or another MIF receptor may play a more vital role.

In seeking to target the interaction of MIF and CD74 on MDSCs we identified Ibudilast as an agent of interest, and were able to treat mice to reduce CD74 expression and increase CD8 T cells in the tumor. One difficulty in using Ibudilast in mouse models is the drug passage effect, where daily treatment increases CYP enzymes leading to rapid degradation^45^. However, in humans the drug is stable in the circulation and accumulates in the CNS with repeated exposer such as daily dosing^44, 52^. For these reasons in the mouse model we settled on a 2x weekly dose of Ibudilast to minimize the drug passage effect, but believe that Ibudilast may be more efficacious in humans than in mouse models. Efforts are currently underway to evaluate Ibudilast in GBM in a clinical trial (NCT03782415)^34^ and will likely provide more insight into how this drug effects the anti-tumor immune response. Additionally, Ibudilast recently demonstrated promising results in a phase 2 clinical trial of multiple sclerosis, where it is thought to have a protective effect by reducing brain atrophy, as compared to anti-inflammatory drugs commonly used to treat multiple sclerosis^35^.

In summary we believe that the M-MDSCs driven by GBM secreted MIF is signaling through the MIF receptor CD74 (**Figure 6**). The interaction between MIF and CD74 subsequently results in pERK signaling, a cascade previously shown in a liver injury model to drive MCP-1 recruiting monocytes to the microenvironment and enhancing the expansion of M-MDSCs (**Figure 6**) ^48, 51, 59^. Our data supports that the signaling pathway initiated by the interaction of MIF and CD74 can be disrupted by the blood brain barrier penetrant small molecule inhibitor, Ibudilast, to reduce M-MDSC function. While our data demonstrates these phenomena, we did not readily observe enhanced survival in our model that involved the use of Ibudilast as a single agent. Nonetheless, we observed that Ibudilast produced an expansion of CD8 T cells and Nanostring analysis predicted an increase multiple pathways including lymphocyte activation. These findings support an interpretation that inhibition of immune suppression, alone, will not be sufficient to produce an anti-tumor immune response. This interpretation mirrors the clinical trial results to date that indicate that treatment with an immune stimulatory therapy alone has been an ineffective strategy. Instead, we hypothesize that better clinical outcomes will be seen when the reversal of tumor-induced immune suppression associated with Ibudilast is combined with an immune stimulatory therapy.

**Figure 6.**
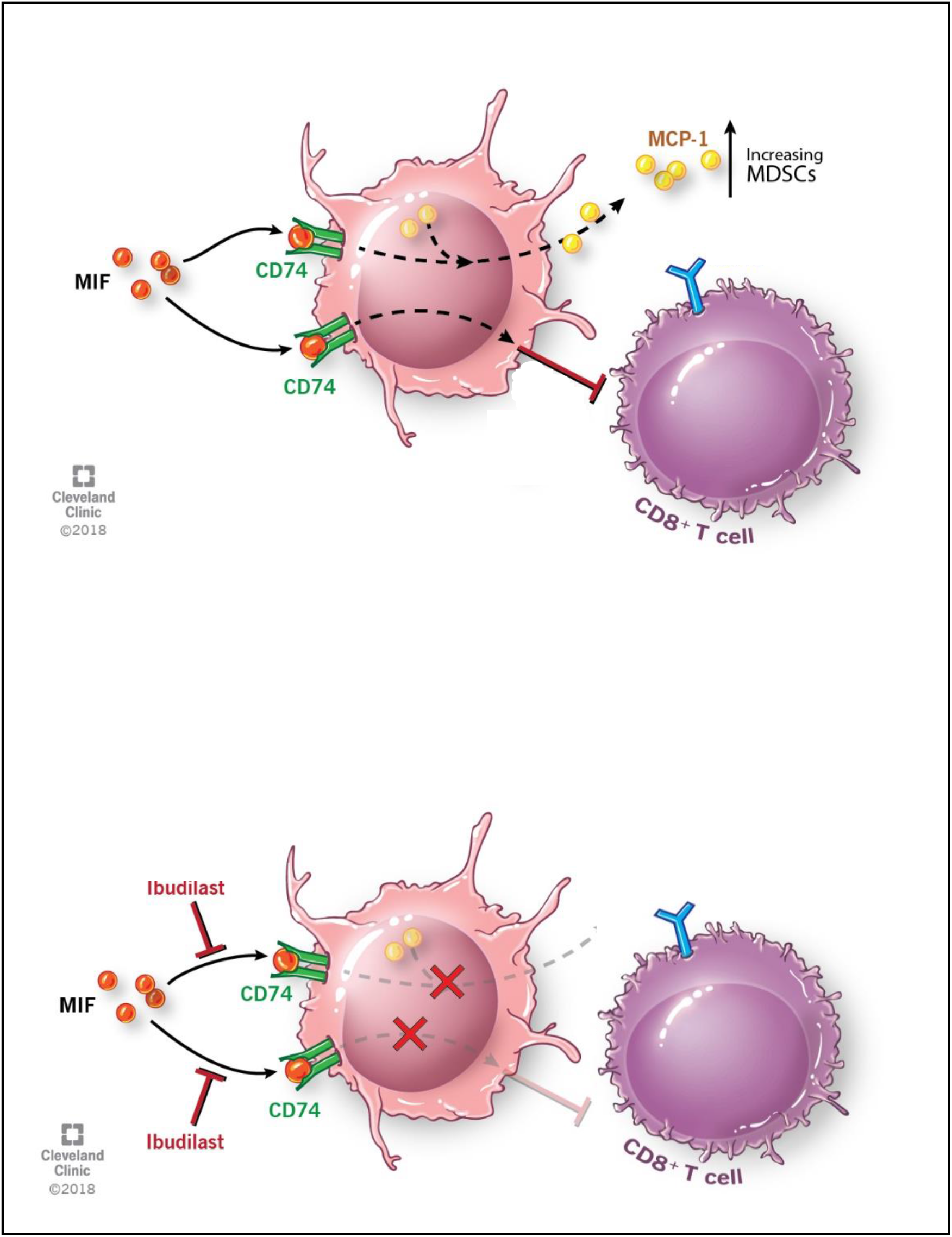
Schematic depicting pathway described where MIF binds CD74 on M-MDSCs enhancing their activity to inhibit CD8 T cells and also produce the downstream target MCP-1. With the addition of Ibudilast to inhibit this process we show a reduction of the MDSCs generation and function removing the inhibitor effect from CD8 T cells.

## Supporting information

Manuscript

## Author Contributions

TJA, DB, BO, KM, KY, RB, MAV and JDL provided conceptualization and design; TJA, DB, AR, GR, LJ, AL, DP, MA, and AM performed the experiments; TJA, DB, BO, LJ, GR, KY, RB, AM, MAV, JDL and analyzed the data; TJA, DB, RB BO, DP, MA, MAV, JDL wrote the manuscript; RB, MAV, and JDL provided financial support; and all authors provided final approval of the manuscript.

## Acknowledgements

We thank the members of the Lathia laboratory for insightful discussion and constructive comments on the manuscript. We thank Joseph Gerow and Eric Schultz for flow cytometry assistance. We thank Amanda Mendelsohn and the Center for Medical Art and Photography at the Cleveland Clinic for providing illustrations and Dr. Erin Mulkearns-Hubert for editorial assistance. This work was funded by an NIH grant (R01 NS109742 to JDL and MAVF31 NS101771 to TJA, F32 CA243314 to DB) the Sontag Foundation (JDL), Blast GBM (JDL, MAV), the Cleveland Clinic VeloSano Bike Race (JDL, MAV), B*CURED (JDL, MAV), the Case Comprehensive Cancer Center (JDL, MAV), and the Cleveland Clinic Brain Tumor Research and Therapeutic Development Research Center of Excellence (MSA, JDL). We also thank Medicnova for supplying Ibudilast for our studies and their collaborative efforts.

## References

1. McGirt MJ, Chaichana KL, Gathinji M, Attenello FJ, Than K, Olivi A, Weingart JD, Brem H and Quinones-Hinojosa AR. Independent association of extent of resection with survival in patients with malignant brain astrocytoma. Journal of neurosurgery. 2009;110:156–62.

2. Stupp R, Hegi ME, Mason WP, van den Bent MJ, Taphoorn MJ, Janzer RC, Ludwin SK, Allgeier A, Fisher B, Belanger K, Hau P, Brandes AA, Gijtenbeek J, Marosi C, Vecht CJ, Mokhtari K, Wesseling P, Villa S, Eisenhauer E, Gorlia T, Weller M, Lacombe D, Cairncross JG, Mirimanoff RO, European Organisation for R, Treatment of Cancer Brain T, Radiation Oncology G and National Cancer Institute of Canada Clinical Trials G. Effects of radiotherapy with concomitant and adjuvant temozolomide versus radiotherapy alone on survival in glioblastoma in a randomised phase III study: 5-year analysis of the EORTC-NCIC trial. The lancet oncology. 2009;10:459–66.

3. Fecci PE, Mitchell DA, Whitesides JF, Xie W, Friedman AH, Archer GE, Herndon JE, 2nd, Bigner DD, Dranoff G and Sampson JH. Increased regulatory T-cell fraction amidst a diminished CD4 compartment explains cellular immune defects in patients with malignant glioma. Cancer research. 2006;66:3294–302.

4. Jacobs JF, Idema AJ, Bol KF, Nierkens S, Grauer OM, Wesseling P, Grotenhuis JA, Hoogerbrugge PM, de Vries IJ and Adema GJ. Regulatory T cells and the PD-L1/PD-1 pathway mediate immune suppression in malignant human brain tumors. Neuro-oncology. 2009;11:394–402.

5. Lewis CE and Pollard JW. Distinct role of macrophages in different tumor microenvironments. Cancer research. 2006;66:605–12.

6. Platten M, Wick W and Weller M. Malignant glioma biology: role for TGF-beta in growth, motility, angiogenesis, and immune escape. Microscopy research and technique. 2001;52:401–10.

7. Huang J, Liu F, Liu Z, Tang H, Wu H, Gong Q and Chen J. Immune Checkpoint in Glioblastoma: Promising and Challenging. Front Pharmacol. 2017;8:242.

8. Heiland DH, Haaker G, Delev D, Mercas B, Masalha W, Heynckes S, Gabelein A, Pfeifer D, Carro MS, Weyerbrock A, Prinz M and Schnell O. Comprehensive analysis of PD-L1 expression in glioblastoma multiforme. Oncotarget. 2017;8:42214–42225.

9. Alban TJ, Alvarado AG, Sorensen MD, Bayik D, Volovetz J, Serbinowski E, Mulkearns-Hubert EE, Sinyuk M, Hale JS, Onzi GR, McGraw M, Huang P, Grabowski MM, Wathen CA, Ahluwalia MS, Radivoyevitch T, Kornblum HI, Kristensen BW, Vogelbaum MA and Lathia JD. Global immune fingerprinting in glioblastoma patient peripheral blood reveals immune-suppression signatures associated with prognosis. JCI Insight. 2018;3.

10. Kamran N, Chandran M, Lowenstein PR and Castro MG. Immature myeloid cells in the tumor microenvironment: Implications for immunotherapy. Clin Immunol. 2018;189:34–42.

11. Kamran N, Kadiyala P, Saxena M, Candolfi M, Li Y, Moreno-Ayala MA, Raja N, Shah D, Lowenstein PR and Castro MG. Immunosuppressive Myeloid Cells’ Blockade in the Glioma Microenvironment Enhances the Efficacy of Immune-Stimulatory Gene Therapy. Mol Ther. 2017;25:232–248.

12. Kohanbash G, McKaveney K, Sakaki M, Ueda R, Mintz AH, Amankulor N, Fujita M, Ohlfest JR and Okada H. GM-CSF promotes the immunosuppressive activity of glioma-infiltrating myeloid cells through interleukin-4 receptor-alpha. Cancer research. 2013;73:6413–23.

13. Raychaudhuri B, Rayman P, Huang P, Grabowski M, Hambardzumyan D, Finke JH and Vogelbaum MA. Myeloid derived suppressor cell infiltration of murine and human gliomas is associated with reduction of tumor infiltrating lymphocytes. J Neurooncol. 2015;122:293–301.

14. Gielen PR, Schulte BM, Kers-Rebel ED, Verrijp K, Petersen-Baltussen HM, ter Laan M, Wesseling P and Adema GJ. Increase in both CD14-positive and CD15-positive myeloid-derived suppressor cell subpopulations in the blood of patients with glioma but predominance of CD15-positive myeloid-derived suppressor cells in glioma tissue. J Neuropathol Exp Neurol. 2015;74:390–400.

15. Chae M, Peterson TE, Balgeman A, Chen S, Zhang L, Renner DN, Johnson AJ and Parney IF. Increasing glioma-associated monocytes leads to increased intratumoral and systemic myeloid-derived suppressor cells in a murine model. Neuro-oncology. 2015;17:978–91.

16. Dubinski D, Wolfer J, Hasselblatt M, Schneider-Hohendorf T, Bogdahn U, Stummer W, Wiendl H and Grauer OM. CD4+ T effector memory cell dysfunction is associated with the accumulation of granulocytic myeloid-derived suppressor cells in glioblastoma patients. Neuro-oncology. 2016;18:807–18.

17. Raychaudhuri B, Rayman P, Ireland J, Ko J, Rini B, Borden EC, Garcia J, Vogelbaum MA and Finke J. Myeloid-derived suppressor cell accumulation and function in patients with newly diagnosed glioblastoma. Neuro-oncology. 2011;13:591–9.

18. Gabrilovich DI and Nagaraj S. Myeloid-derived suppressor cells as regulators of the immune system. Nat Rev Immunol. 2009;9:162–74.

19. Gielen PR, Schulte BM, Kers-Rebel ED, Verrijp K, Bossman SA, Ter Laan M, Wesseling P and Adema GJ. Elevated levels of polymorphonuclear myeloid-derived suppressor cells in patients with glioblastoma highly express S100A8/9 and arginase and suppress T cell function. Neuro-oncology. 2016;18:1253–64.

20. Peereboom DM, Alban TJ, Grabowski MM, Alvarado AG, Otvos B, Bayik D, Roversi G, McGraw M, Huang P, Mohammadi AM, Kornblum HI, Radivoyevitch T, Ahluwalia MS, Vogelbaum MA and Lathia JD. Metronomic capecitabine as an immune modulator in glioblastoma patients reduces myeloid-derived suppressor cells. JCI Insight. 2019;4.

21. Otvos B, Silver DJ, Mulkearns-Hubert EE, Alvarado AG, Turaga SM, Sorensen MD, Rayman P, Flavahan WA, Hale JS, Stoltz K, Sinyuk M, Wu Q, Jarrar A, Kim SH, Fox PL, Nakano I, Rich JN, Ransohoff RM, Finke J, Kristensen BW, Vogelbaum MA and Lathia JD. Cancer stem cell-secreted macrophage migration inhibitory factor stimulates myeloid derived suppressor cell function and facilitates glioblastoma immune evasion. Stem Cells. 2016;34:2026–39.

22. Simpson KD, Templeton DJ and Cross JV. Macrophage migration inhibitory factor promotes tumor growth and metastasis by inducing myeloid-derived suppressor cells in the tumor microenvironment. J Immunol. 2012;189:5533–40.

23. Castro BA, Flanigan P, Jahangiri A, Hoffman D, Chen W, Kuang R, De Lay M, Yagnik G, Wagner JR, Mascharak S, Sidorov M, Shrivastav S, Kohanbash G, Okada H and Aghi MK. Macrophage migration inhibitory factor downregulation: a novel mechanism of resistance to anti-angiogenic therapy. Oncogene. 2017;36:3749–3759.

24. Das R, Koo MS, Kim BH, Jacob ST, Subbian S, Yao J, Leng L, Levy R, Murchison C, Burman WJ, Moore CC, Scheld WM, David JR, Kaplan G, MacMicking JD and Bucala R. Macrophage migration inhibitory factor (MIF) is a critical mediator of the innate immune response to Mycobacterium tuberculosis. Proceedings of the National Academy of Sciences of the United States of America. 2013;110:E2997–3006.

25. Calandra T and Roger T. Macrophage migration inhibitory factor: a regulator of innate immunity. Nat Rev Immunol. 2003;3:791–800.

26. Conroy H, Mawhinney L and Donnelly SC. Inflammation and cancer: macrophage migration inhibitory factor (MIF)--the potential missing link. QJM. 2010;103:831–6.

27. Hoi AY, Iskander MN and Morand EF. Macrophage migration inhibitory factor: a therapeutic target across inflammatory diseases. Inflamm Allergy Drug Targets. 2007;6:183–90.

28. Yaddanapudi K, Rendon BE, Lamont G, Kim EJ, Al Rayyan N, Richie J, Albeituni S, Waigel S, Wise A and Mitchell RA. MIF Is Necessary for Late-Stage Melanoma Patient MDSC Immune Suppression and Differentiation. Cancer Immunol Res. 2016;4:101–12.

29. Figueiredo CR, Azevedo RA, Mousdell S, Resende-Lara PT, Ireland L, Santos A, Girola N, Cunha R, Schmid MC, Polonelli L, Travassos LR and Mielgo A. Blockade of MIF-CD74 Signalling on Macrophages and Dendritic Cells Restores the Antitumour Immune Response Against Metastatic Melanoma. Front Immunol. 2018;9:1132.

30. Subbannayya T, Variar P, Advani J, Nair B, Shankar S, Gowda H, Saussez S, Chatterjee A and Prasad TS. An integrated signal transduction network of macrophage migration inhibitory factor. J Cell Commun Signal. 2016;10:165–70.

31. Jankauskas SS, Wong DWL, Bucala R, Djudjaj S and Boor P. Evolving complexity of MIF signaling. Cell Signal. 2019;57:76–88.

32. Shi X, Leng L, Wang T, Wang W, Du X, Li J, McDonald C, Chen Z, Murphy JW, Lolis E, Noble P, Knudson W and Bucala R. CD44 is the signaling component of the macrophage migration inhibitory factor-CD74 receptor complex. Immunity. 2006;25:595–606.

33. O’Reilly C, Doroudian M, Mawhinney L and Donnelly SC. Targeting MIF in Cancer: Therapeutic Strategies, Current Developments, and Future Opportunities. Med Res Rev. 2016;36:440–60.

34. Ha W, Sevim-Nalkiran H, Zaman AM, Matsuda K, Khasraw M, Nowak AK, Chung L, Baxter RC and McDonald KL. Ibudilast sensitizes glioblastoma to temozolomide by targeting Macrophage Migration Inhibitory Factor (MIF). Sci Rep. 2019;9:2905.

35. Fox RJ, Coffey CS, Conwit R, Cudkowicz ME, Gleason T, Goodman A, Klawiter EC, Matsuda K, McGovern M, Naismith RT, Ashokkumar A, Barnes J, Ecklund D, Klingner E, Koepp M, Long JD, Natarajan S, Thornell B, Yankey J, Bermel RA, Debbins JP, Huang X, Jagodnik P, Lowe MJ, Nakamura K, Narayanan S, Sakaie KE, Thoomukuntla B, Zhou X, Krieger S, Alvarez E, Apperson M, Bashir K, Cohen BA, Coyle PK, Delgado S, Dewitt LD, Flores A, Giesser BS, Goldman MD, Jubelt B, Lava N, Lynch SG, Moses H, Ontaneda D, Perumal JS, Racke M, Repovic P, Riley CS, Severson C, Shinnar S, Suski V, Weinstock-Guttman B, Yadav V, Zabeti A and Investigators NS-MT. Phase 2 Trial of Ibudilast in Progressive Multiple Sclerosis. N Engl J Med. 2018;379:846–855.

36. Kim KW and Kim HR. Macrophage migration inhibitory factor: a potential therapeutic target for rheumatoid arthritis. Korean J Intern Med. 2016;31:634–42.

37. Bilsborrow JB, Doherty E, Tilstam PV and Bucala R. Macrophage migration inhibitory factor (MIF) as a therapeutic target for rheumatoid arthritis and systemic lupus erythematosus. Expert Opin Ther Targets. 2019;23:733–744.

38. Oliver J, Marquez A, Gomez-Garcia M, Martinez A, Mendoza JL, Vilchez JR, Lopez-Nevot MA, Pinero A, de la Concha EG, Nieto A, Urcelay E and Martin J. Association of the macrophage migration inhibitory factor gene polymorphisms with inflammatory bowel disease. Gut. 2007;56:150–1.

39. Yang H, Zheng S, Mao Y, Chen Z, Zheng C, Li H, Sumners C, Li Q, Yang P and Lei B. Modulating of ocular inflammation with macrophage migration inhibitory factor is associated with notch signalling in experimental autoimmune uveitis. Clin Exp Immunol. 2016;183:280–93.

40. Simpson KD and Cross JV. MIF: metastasis/MDSC-inducing factor? Oncoimmunology. 2013;2:e23337.

41. Bronte V, Brandau S, Chen SH, Colombo MP, Frey AB, Greten TF, Mandruzzato S, Murray PJ, Ochoa A, Ostrand-Rosenberg S, Rodriguez PC, Sica A, Umansky V, Vonderheide RH and Gabrilovich DI. Recommendations for myeloid-derived suppressor cell nomenclature and characterization standards. Nat Commun. 2016;7:12150.

42. Darmanis S, Sloan SA, Croote D, Mignardi M, Chernikova S, Samghababi P, Zhang Y, Neff N, Kowarsky M, Caneda C, Li G, Chang SD, Connolly ID, Li Y, Barres BA, Gephart MH and Quake SR. Single-Cell RNA-Seq Analysis of Infiltrating Neoplastic Cells at the Migrating Front of Human Glioblastoma. Cell Rep. 2017;21:1399–1410.

43. Cournia Z, Leng L, Gandavadi S, Du X, Bucala R and Jorgensen WL. Discovery of human macrophage migration inhibitory factor (MIF)-CD74 antagonists via virtual screening. J Med Chem. 2009;52:416–24.

44. Rolan P, Hutchinson M and Johnson K. Ibudilast: a review of its pharmacology, efficacy and safety in respiratory and neurological disease. Expert Opin Pharmacother. 2009;10:2897–904.

45. Sanftner LM, Gibbons JA, Gross MI, Suzuki BM, Gaeta FC and Johnson KW. Cross-species comparisons of the pharmacokinetics of ibudilast. Xenobiotica. 2009;39:964–77.

46. Cho Y, Crichlow GV, Vermeire JJ, Leng L, Du X, Hodsdon ME, Bucala R, Cappello M, Gross M, Gaeta F, Johnson K and Lolis EJ. Allosteric inhibition of macrophage migration inhibitory factor revealed by ibudilast. Proceedings of the National Academy of Sciences of the United States of America. 2010;107:11313–8.

47. Yoo SA, Leng L, Kim BJ, Du X, Tilstam PV, Kim KH, Kong JS, Yoon HJ, Liu A, Wang T, Song Y, Sauler M, Bernhagen J, Ritchlin CT, Lee P, Cho CS, Kim WU and Bucala R. MIF allele-dependent regulation of the MIF coreceptor CD44 and role in rheumatoid arthritis. Proceedings of the National Academy of Sciences of the United States of America. 2016;113:E7917–E7926.

48. Leng L, Chen L, Fan J, Greven D, Arjona A, Du X, Austin D, Kashgarian M, Yin Z, Huang XR, Lan HY, Lolis E, Nikolic-Paterson D and Bucala R. A small-molecule macrophage migration inhibitory factor antagonist protects against glomerulonephritis in lupus-prone NZB/NZW F1 and MRL/lpr mice. J Immunol. 2011;186:527–38.

49. Hare AA, Leng L, Gandavadi S, Du X, Cournia Z, Bucala R and Jorgensen WL. Optimization of N-benzyl-benzoxazol-2-ones as receptor antagonists of macrophage migration inhibitory factor (MIF). Bioorg Med Chem Lett. 2010;20:5811–4.

50. Najjar YG and Finke JH. Clinical perspectives on targeting of myeloid derived suppressor cells in the treatment of cancer. Front Oncol. 2013;3:49.

51. Xie J, Yang L, Tian L, Li W, Yang L and Li L. Macrophage Migration Inhibitor Factor Upregulates MCP-1 Expression in an Autocrine Manner in Hepatocytes during Acute Mouse Liver Injury. Sci Rep. 2016;6:27665.

52. Rolan P, Gibbons JA, He L, Chang E, Jones D, Gross MI, Davidson JB, Sanftner LM and Johnson KW. Ibudilast in healthy volunteers: safety, tolerability and pharmacokinetics with single and multiple doses. Br J Clin Pharmacol. 2008;66:792–801.

53. Schwenkgrub J, Zaremba M, Mirowska-Guzel D and Kurkowska-Jastrzebska I. Ibudilast: a nonselective phosphodiesterase inhibitor in brain disorders. Postepy Hig Med Dosw (Online). 2017;71:137–148.

54. Schwenkgrub J, Zaremba M, Joniec-Maciejak I, Cudna A, Mirowska-Guzel D and Kurkowska-Jastrzebska I. The phosphodiesterase inhibitor, ibudilast, attenuates neuroinflammation in the MPTP model of Parkinson’s disease. PLoS One. 2017;12:e0182019.

55. MacKenzie SJ and Houslay MD. Action of rolipram on specific PDE4 cAMP phosphodiesterase isoforms and on the phosphorylation of cAMP-response-element-binding protein (CREB) and p38 mitogen-activated protein (MAP) kinase in U937 monocytic cells. Biochem J. 2000;347:571–8.

56. Zhang H, Ye YL, Li MX, Ye SB, Huang WR, Cai TT, He J, Peng JY, Duan TH, Cui J, Zhang XS, Zhou FJ, Wang RF and Li J. CXCL2/MIF-CXCR2 signaling promotes the recruitment of myeloid-derived suppressor cells and is correlated with prognosis in bladder cancer. Oncogene. 2017;36:2095–2104.

57. Zilio S and Serafini P. Neutrophils and Granulocytic MDSC: The Janus God of Cancer Immunotherapy. Vaccines (Basel). 2016;4.

58. Clavijo PE, Moore EC, Chen J, Davis RJ, Friedman J, Kim Y, Van Waes C, Chen Z and Allen CT. Resistance to CTLA-4 checkpoint inhibition reversed through selective elimination of granulocytic myeloid cells. Oncotarget. 2017;8:55804–55820.

59. Hoi AY, Hickey MJ, Hall P, Yamana J, O’Sullivan KM, Santos LL, James WG, Kitching AR and Morand EF. Macrophage migration inhibitory factor deficiency attenuates macrophage recruitment, glomerulonephritis, and lethality in MRL/lpr mice. J Immunol. 2006;177:5687–96.

