## Supplementary material for "Glioblastoma myeloid-derived suppressor cell subsets express differential macrophage migration inhibitory factor receptor profiles that can be targeted to reduce immune suppression": Manuscript

Supplemental Figure 1. MIF knockdown reduced myeloid infiltration into tumors and does not increase survival in a mouse mode of GBM using immune incompetent mice

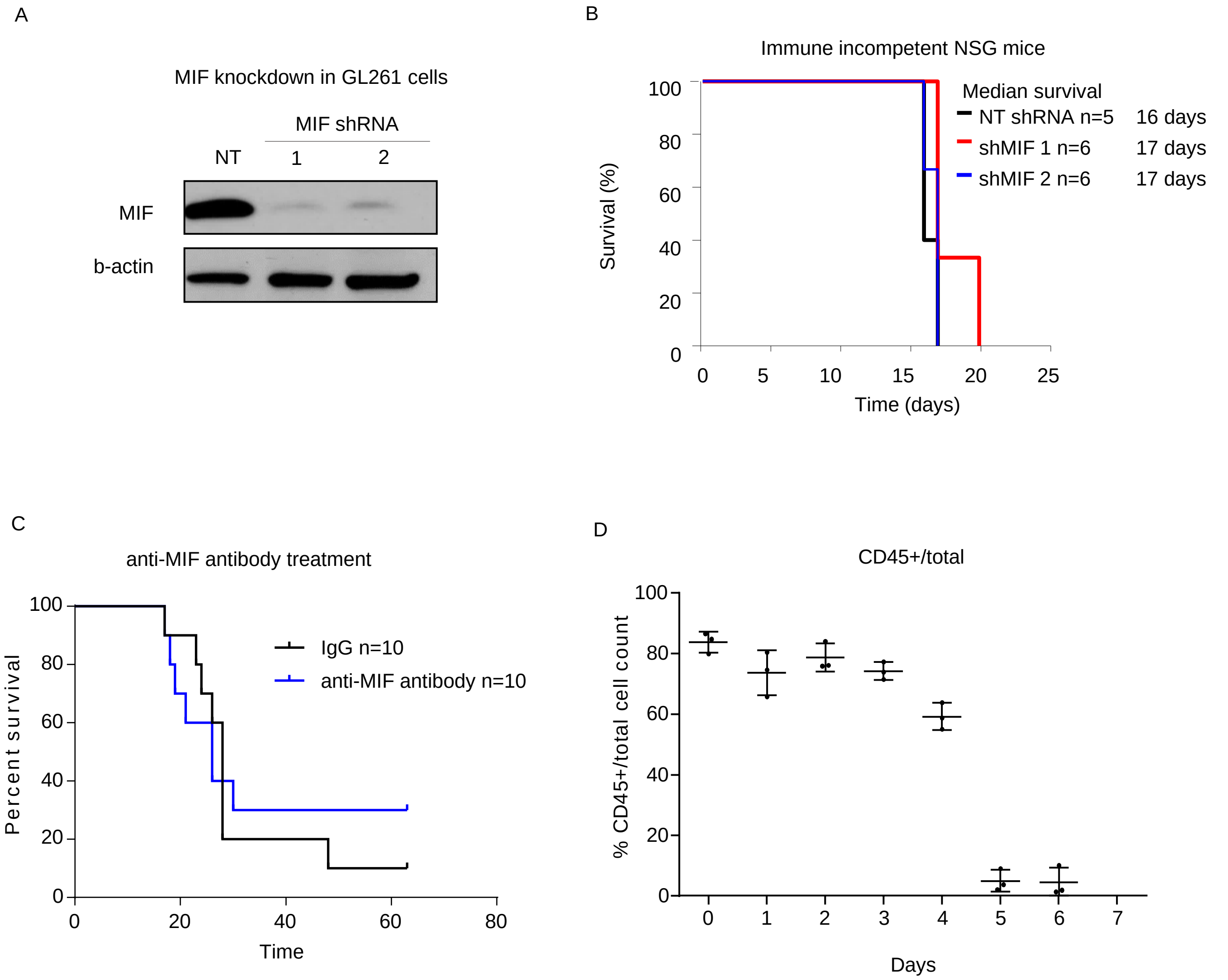

Supplemental Figure 2.

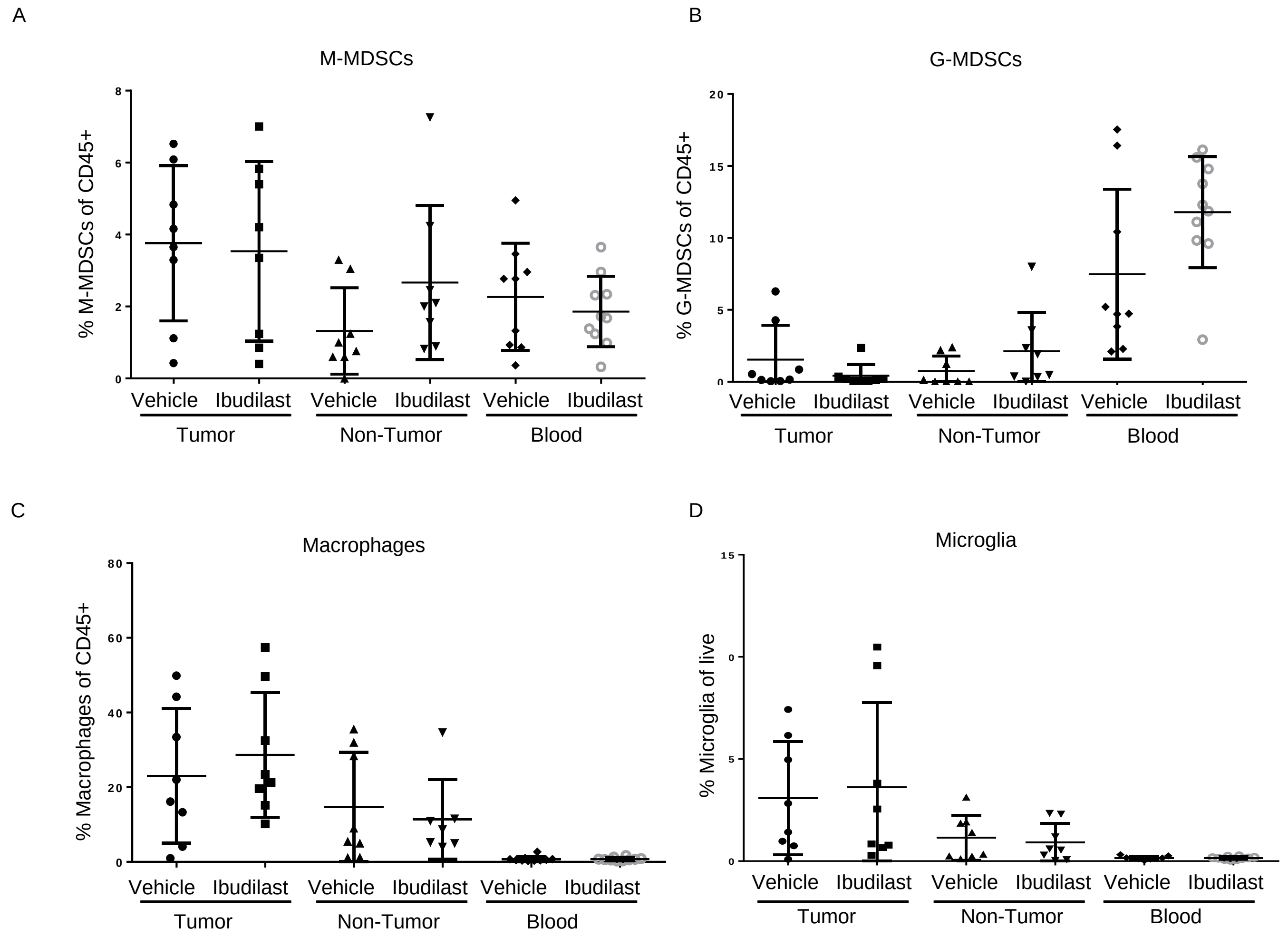

Supplemental Figure 3.

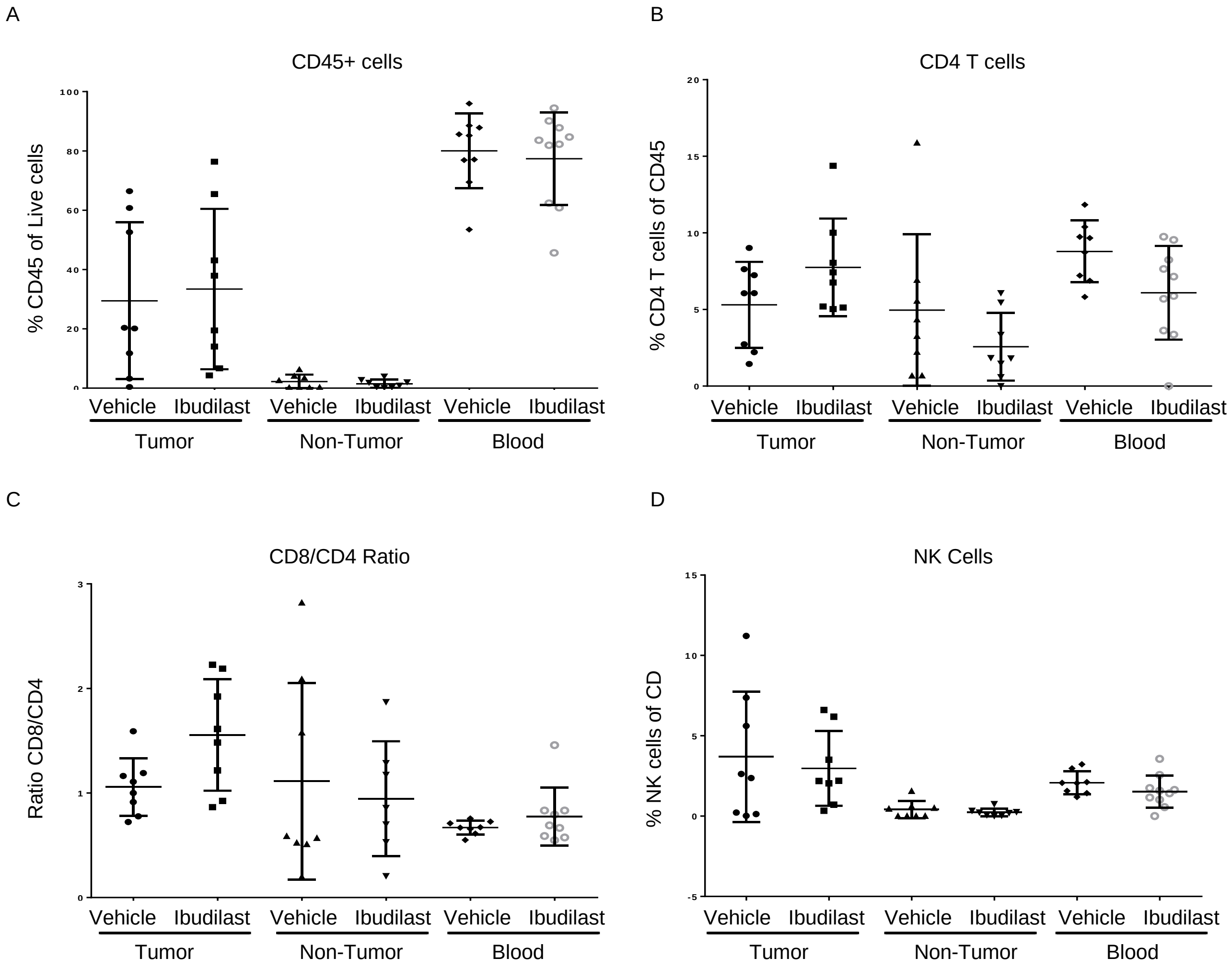

Supplemental Figure 4.

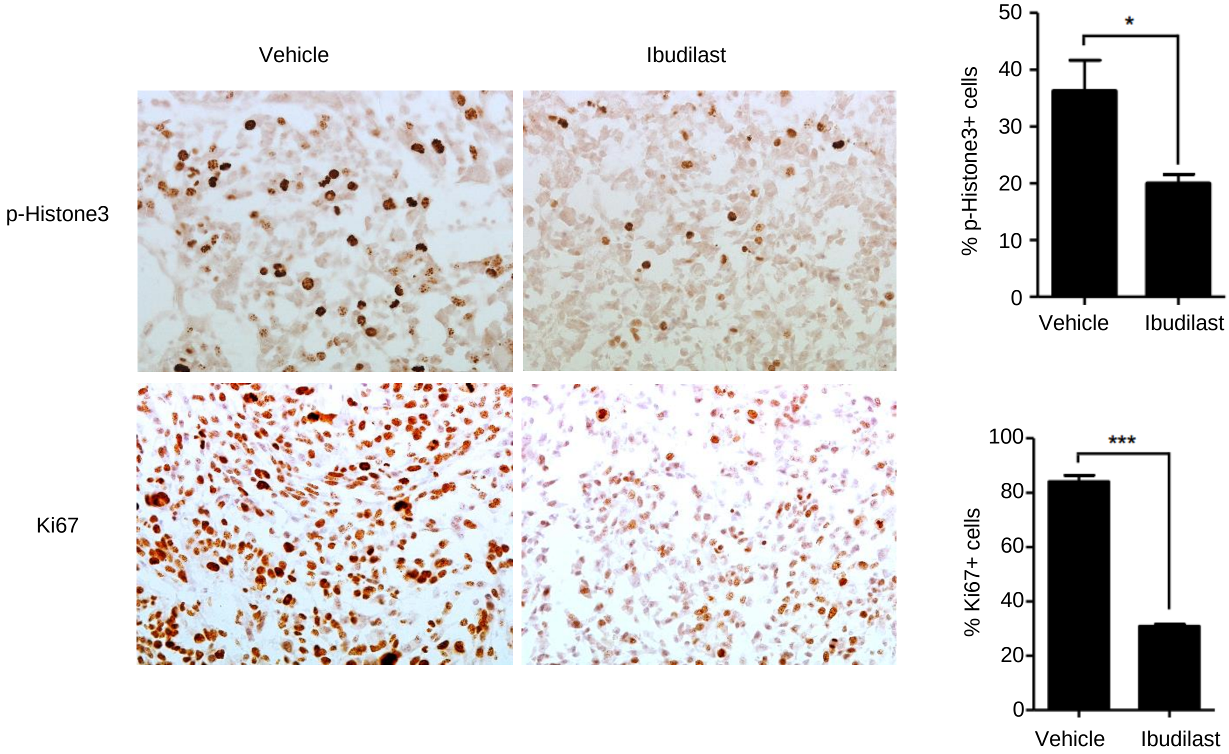

**Supplemental Figure 1.** shRNA knockdown of MIF in GL261 was performed using 2 separate shRNA's which were the top targets from previously published work from our group to generate stable knockdown cell lines of GL261 (**A**). Comparing the survival of intracranially implanted tumors in NSG immune incompetent mice demonstrate no survival difference in NSG mice (**B**). Treating n=10 GL261 tumor bearing mice 2x weekly with anti-MIF antibody (gifted from Dr. Richard Bucala) vs n=10 IgG control treated mice demonstrated no survival benefit (**C**). MDSC co-culture dynamics over time analyzing n=3 mice in separate co-cultures where one well was used each day over 7 days to check the number of CD45+ cells by flow cytometry (**D**). Survival curve analysis was performed in GraphPad Prism using Log-rank (Mantel-Cox) test for p value and hazard ratio log rank was computed on the same data using GraphPad Prism.

**Supplemental Figure 2.** Intracrainially injected tumors vehicle vs ibudilast treated tumors, non-tumor tissue, and blood analysis from **Figure 5** demonstrate no significant difference in M-MDSCs, G-MDSCs, Macrophages, or Microglia (**A-D**). Two-Tailed T-Test was performed for comparing vehicle vs ibudilast in each compartment *<0.05, ** <0.01, ***<0.001.

**Supplemental Figure 3.** Intracrainially injected tumors vehicle vs ibudilast treated tumors, non-tumor tissue, and blood analysis from **Figure 5** demonstrate no significant difference in total CD45+ cells, CD4 T cells, ratio of CD8/ CD4 T cells, or NK cells (**A-D**). Two-Tailed T-Test was performed for comparing vehicle vs ibudilast in each compartment *<0.05, ** <0.01, ***<0.001.

**Supplemental Figure 4.** Intracrainially injected tumors vehicle vs ibudilast treated mice were perfused at endpoint and tissue was paraffin embedded for IHC analysis. Staining for p-Histone3 and Ki67 demonstrated a reduction in proliferation in the ibudilast treated tumors. Two-Tailed T-Test was performed for comparing vehicle (mock) vs ibudilast treated tumors *<0.05, ** <0.01, ***<0.001
